# Selection and gene flow shape niche-associated copy-number variation of pheromone receptor genes

**DOI:** 10.1101/580803

**Authors:** Daehan Lee, Stefan Zdraljevic, Daniel E. Cook, Lise Frézal, Jung-Chen Hsu, Mark G. Sterken, Joost A.G. Riksen, John Wang, Jan E. Kammenga, Christian Braendle, Marie-Anne Félix, Frank C. Schroeder, Erik C. Andersen

## Abstract

From quorum sensing in bacteria to pheromone signaling in social insects, chemical communication mediates interactions among individuals in a local population. In *Caenorhabditis elegans*, ascaroside pheromones can dictate local population density, in which high levels of pheromones inhibit the reproductive maturation of individuals. Little is known about how natural genetic diversity affects the pheromone responses of individuals from diverse habitats. Here, we show that a niche-associated copy-number variation (CNV) of pheromone receptor genes contributes to natural differences in pheromone responses. We found putative loss-of-function deletions that reduce copy number of duplicated pheromone receptor genes (*srg-36 and srg-37*), which were shown previously to be selected in population-dense laboratory cultures. A common natural deletion in the less functional copy (*srg-37*) arose from a single ancestral population that spread throughout the world and underlies reduced pheromone sensitivity across the global *C. elegans* population. This deletion is enriched in wild strains that were isolated from a rotting fruit niche, where proliferating populations are often found. Taken together, these results demonstrate that selection and gene flow together shape the copy number of pheromone receptor genes in natural *C. elegans* populations to facilitate local adaptation to diverse niches.

## Introduction

To maximize reproductive success, organisms must respond to changing environmental conditions. In a fluctuating environment, each response will likely have a fitness trade-off with reproductive success now or in the future. *Caenorhabditis elegans* can either grow to a reproductive adult in three days or delay maturity for months by entering the dauer diapause stage^1^. Food supply and pheromone signal are two major inputs that affect this developmental plasticity^2^. The accumulation of ascaroside pheromones in dense populations antagonizes food signals that promote growth and inhibits the induction of the stress-resistant and long-lived dauer stage^3, 4^. Animals must measure the amount of remaining food and the pheromone concentration to determine if it is advantageous to continue reproductive growth or enter the dauer stage to disperse and hopefully encounter a new food source. Therefore, the decision to enter dauer decreases reproductive success in the short-term in favor of future survival success. Decades of research have provided insights into the chemical and genetic basis of the dauer-pheromone response^5^. However, most studies used a single laboratory-adapted strain (N2), which has limited our understanding of the natural processes that have shaped the dauer-pheromone response.

After decades of focused laboratory research on *C. elegans* as a model organism, the natural history of this species has only recently been described from extensive field research^6^. These field studies have revealed that the dauer stage is important for the population dynamics in their natural habitat^7^. These dynamics are typified by a “boom” phase after initial colonization of a nutrient-rich habitat, followed by a “bust” phase when resources are depleted. At the end of the boom phase when the local population size is large and nutrients are limited, individual animals enter the dauer stage. Dauers exhibit a stage-specific behavior called nictation, which facilitates phoretic interactions between dauer larvae and more mobile animals to disperse to favorable environments^8, 9^. Because dauer larvae are presumed to play a crucial role in the survival and dispersal of the species, it is likely that the decision to enter the dauer stage is under natural selection. Although differences in dauer development among small number of wild *C. elegans* strains have been described previously^10–15^, no underlying natural genetic variant has been identified yet. Here, we integrate laboratory experiments, computational genomic analyses, and field research to further our understanding of the genetic basis underlying microevolution of the pheromone-mediated developmental plasticity. We identify the natural genetic variation of dauer-pheromone responses and characterize a copy-number variation of pheromone receptor genes that has been shaped by niche-associated selection and gene flow.

## RESULTS

### Natural variation of the dauer-pheromone response was measured using a high-throughput dauer assay

To explore the effects of natural genetic variation on the ability to enter the dauer stage, we developed a high-throughput assay (HTDA) to quantify the dauer-pheromone responses of wild *C. elegans* strains. The HTDA takes advantage of the observation that dauer larvae have no pharyngeal pumping^16^. We treated animals with fluorescent microspheres that can be ingested and then quantified both fluorescence and size of individual animals using a large-particle flow cytometer (COPAS BIOSORT, Union Biometrica). These data facilitated computational classification of dauers (Fig. 1a, b; Materials and methods) and recapitulated the known differences in the dauer-pheromone responses between N2 and a constitutive dauer mutant *daf-2(e1370)*, as well as the dauer-inducing effect of synthetic pheromone (Fig. 1b, c). To determine if genetic variation within the *C. elegans* species causes differential dauer-pheromone responses, we applied the HTDA to four genetically divergent *C. elegans* strains after treatment with various concentrations of four known dauer-inducing synthetic ascarosides (ascr#2, ascr#3, ascr#5, and ascr#8). We found significant variation in the dauer-pheromone response among the strains tested, as measured by the fraction of individuals that enter the dauer stage (Fig. 1d, Supplementary Fig. 1). Among the conditions we tested, we found that 800 nM ascr#5 maximizes the among-strain variance and minimizes the within-strain variance in dauer-pheromone response. These results enabled us to survey the effects of genetic variation on the dauer-pheromone response across the *C. elegans* species.

**Fig. 1:**
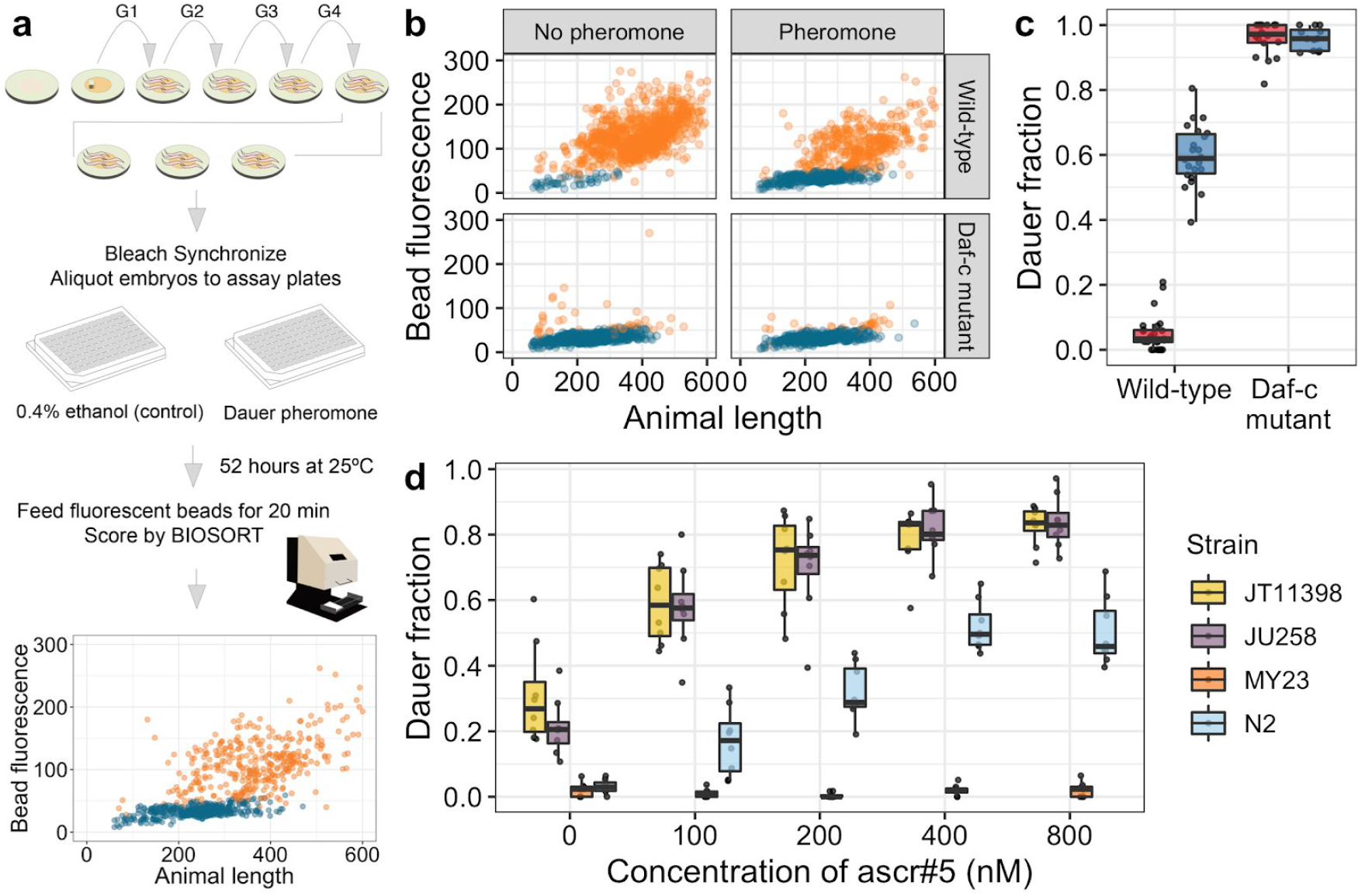
A high-throughput dauer assay measures natural variation of dauer-pheromone response. (a) The workflow for the high-throughput dauer assay (HTDA) using a COPAS BIOSORT is shown (see Materials and methods for further description). (b) Optical measurements of the laboratory wild-type strain (N2) (Top) and a Daf-c mutant *daf-2(e1370)* (bottom) are shown under control (left) and pheromone-treated (ascr#5 800 nM) conditions (right) at 25°C using the HTDA. Animal size and fluorescence-intensity traits are used as variables to build a model that differentiates dauer (blue) and non-dauer populations (orange). Relative animal length measured by time-of-flight (TOF) is shown on the x-axis, and bead-derived fluorescent intensity is shown on the y-axis. (c) Tukey box plots of the dauer fraction quantification from (b) are shown with data points plotted behind. Box plots are colored by assay conditions (control (red) and ascr#5 800 nM treatment (blue)). The genotypes are shown on the x-axis, and fractions of dauer larvae are shown on the y-axis. (d) Tukey box plots of the ascr#5 dose response at 25°C for four divergent strains are shown with data points plotted behind. Box plots are colored by strain (JT11398 (yellow), JU258 (purple), MY23 (orange) and N2 (blue)). Concentrations of ascr#5 are shown on the x-axis, and fractions of dauer larvae are shown on the y-axis.

### Genome-wide association (GWA) mapping reveals multiple loci underlying natural variation of the ascr#5 response

Next, we quantified dauer induction of 157 wild strains that have been isolated from diverse habitats across six continents (Supplementary Fig. 2)^17, 18^. We found significant variation in the ascr#5 response with high broad-sense heritability (0.83), indicating that most of the observed phenotypic variance can be explained by genetic differences across these strains (Fig. 2a). The two strains that represent the phenotypic extremes of the ascr#5 response are EG4349 and JU2576, where EG4349 did not enter dauer and was completely insensitive to ascr#5 treatment, and a large fraction of the JU2576 individuals entered the dauer stage in the same condition. Overall, we observed a continuous distribution of dauer-pheromone responses among these wild strains (mean = 0.41, standard deviation = 0.20), indicating that natural variation in this trait is likely not explained by a single gene.

**Fig. 2:**
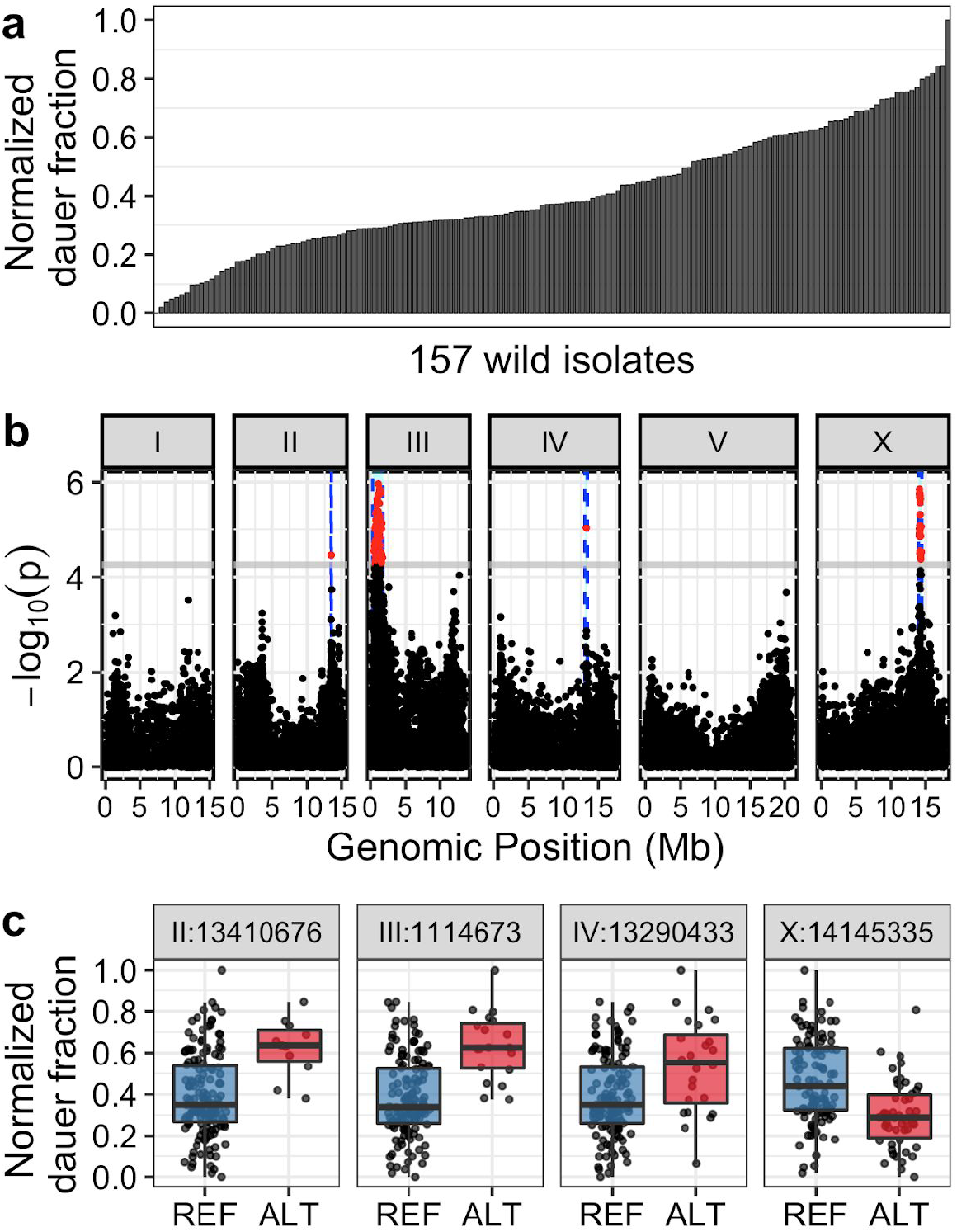
Genome-wide association (GWA) mapping reveals four major loci underlying natural variation in dauer-pheromone response. (a) A bar plot for the natural variation of ascr#5-induced dauer formation at 25°C across 157 *C. elegans* wild isolates (one-way analysis of variance (ANOVA), log_10_*p* = −49.6598) is shown. Each bar represents the phenotypic response of a single wild isolate to 800 nM ascr#5. (b) A manhattan plot for single-marker based GWA mapping of the ascr#5-induced dauer formation trait from (a) is shown. Each dot represents a single-nucleotide variant (SNV) that is present in at least 5% of the 249 wild strains. The genomic position in Mb, separated by chromosome, is plotted on the x-axis, and the statistical significance of the correlation between genotype and phenotype is plotted on the y-axis. SNVs are colored red if they pass the genome-wide eigen-decomposition significance threshold, which is denoted by the gray horizontal line. The region of interest for each QTL is represented by vertical blue dotted lines. (c) Tukey box plots of phenotypes split by peak marker position of the four QTL (chrII:13410676, chrIII:1114673, chrIV:13290433, chrX:14145335) are shown. Each dot corresponds to the phenotype of an individual strain, which is plotted on the y-axis as the normalized dauer fraction phenotype. Strains are grouped by their genotype at each peak QTL position, where REF (blue) corresponds to the reference allele from the laboratory N2 strain and ALT (red) corresponds to the alternative allele.

To characterize the quantitative trait loci (QTL) associated with variation in the ascr#5 response, we performed genome-wide association (GWA) mappings and identified four QTL (Fig. 2b, c). The QTL that explained the most variation in pheromone-induced dauer induction (15.9%) is on the right arm of the X chromosome. Strains that have the non-reference (ALT) allele at the peak marker (X:14,145,335) of this QTL were less responsive to ascr#5 treatment than strains that have the reference (REF) allele (REF mean: 0.46; ALT mean: 0.30, log_10_*p* = −5.851505). The remaining QTL on chromosomes II, III, and IV, explain 8.4%, 15.1%, and 5.4% of the variation in the ascr#5 response, respectively. We also found no obvious linkage disequilibrium (LD) among these QTL (Supplementary Fig. 3), suggesting that multiple genomic loci underlie natural variation in the ascr#5 response.

### A putative loss-of-function allele in an ascr#5 receptor gene is associated with reduced dauer formation

We focused our efforts on the largest effect QTL, which we named *dauf-1* (dauer-formation QTL #1). The 469-kb surrounding the *dauf-1* peak marker contains 82 protein-coding genes, including the duplicated genes *srg-36* and *srg-37*, which encode ascr#5 receptors^19^. Both genes are expressed in the same pair of chemosensory neurons (ASI), which play an essential role in the dauer-pheromone response^20, 21^. Notably, previous studies reported that these two genes are repeatedly eliminated during long-term propagation of *C. elegans* in high-density liquid cultures^19^. Large deletions that remove both *srg-36* and *srg-37* arose in two independent laboratory-domesticated lineages, causing the loss of sensitivity to ascr#5.

To evaluate whether similar mutations in these two genes underlie the *dauf-1* QTL, we investigated the genome sequences of 249 wild strains available through the *C. elegans* Natural Diversity Resource (CeNDR)^22, 23^. Although we could not find a large deletion that removes both *srg-36* and *srg-37*, we found one strain with a 411-bp deletion in *srg-36* and many other strains with an identical 94-bp deletion in *srg-37* (Fig. 3a, Supplementary Fig. 4). We named these deletions *srg-36(ean178)* and *srg-37(ean179)*. To test whether these deletions can explain the *dauf-1* QTL effect, we analyzed the association between the ascr#5 response and the two deletions. First, we found that *srg-36(ean178)*, which is a deletion found only in the PB303 strain and removes the fourth and fifth exons, is associated with an insensitivity to a high dose of ascr#5 (2 µM) (Supplementary Fig. 5). Because this deletion allele was not found in any other wild strains, *srg-36(ean178)* cannot explain the population-wide differences in dauer formation. By contrast, we found that all wild strains with the *srg-37(ean179)* deletion belong to *dauf-1(ALT)* group and had reduced ascr#5 sensitivity (Fig. 3b), suggesting that this deletion allele might cause a reduction in the acr#5 response.

**Fig. 3:**
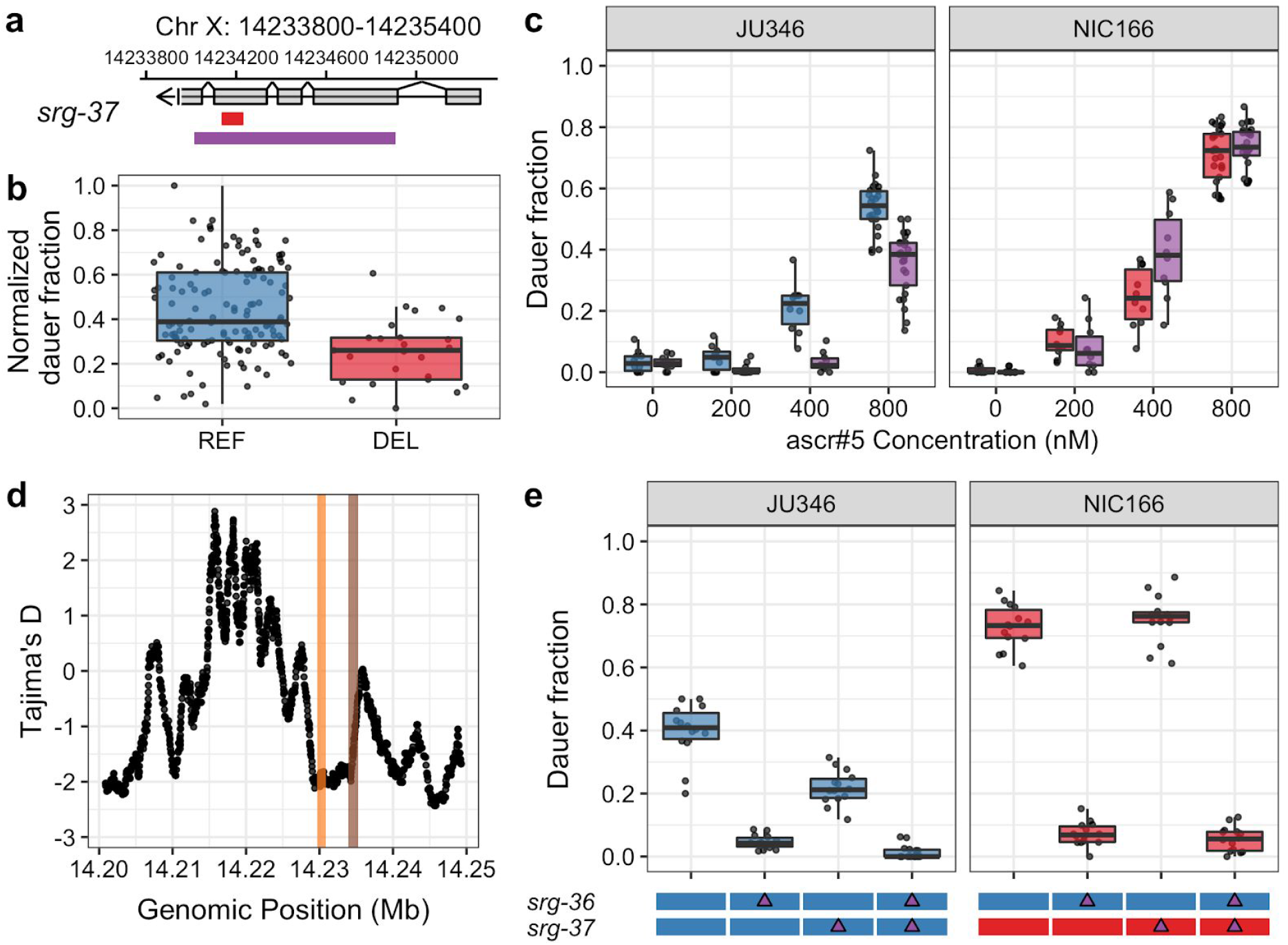
Natural variant in the ascr#5 receptor gene, *srg-37*, underlies natural differences in dauer formation. (a) A schematic plot for the *srg-37* gene structure (grey), 94-bp natural deletion allele *ean179* (red), and CRISPR-Cas9 genome-editing target sequences for the putative loss-of-function deletion (purple) are shown. (b) Tukey box plots of dauer formation split by *srg-37* genotype are shown. Each dot corresponds to the phenotype of an individual strain, which is plotted on the y-axis by the normalized dauer fraction. Strains are grouped by their *srg-37* genotype, where REF (blue) corresponds to the wild-type reference allele from the laboratory N2 strain and DEL (red) corresponds to the natural 94-bp deletion allele (*ean179*). (c) Tukey box plots of the ascr#5 dose-response differences at 25°C among two wild isolates and *srg-37(lf)* mutants in both backgrounds are shown with data points plotted behind. A dose response comparison is shown between (Left) JU346 *srg-37(+)* (blue) and JU346 *srg-37(lf)* (purple) and (Right) NIC166 *srg-37(ean179)* (red) and NIC166 *srg-37(lf)* (purple). The concentration of ascr#5 is shown on the x-axis, and the fraction of dauer formation is shown on the y-axis. (d) Tajima’s D statistics across the *srg-36 srg-37* locus are shown. Each dot corresponds to a Tajima’s D statistic calculated from the allele frequency spectrum of 50 SNVs across 249 wild isolates. Colored lines highlight coding regions of *srg-36* (orange) and *srg-37* (brown). (e) Tukey box plots of *srg-36* and *srg-37* loss-of-function experiments under control (red, 0.4% ethanol) and ascr#5 pheromone conditions (blue, 2 µM of ascr#5) at 25°C are shown with data points plotted behind. Genotypes of *srg-36* and *srg-37* are shown as colored bars on the x-axis, where red bar corresponds to the genotype with the natural deletion allele, *srg-37(ean179)*, and the blue bar corresponds to the genotype without the deletion. Purple triangles represent the CRISPR-Cas9-mediated loss-of-function mutation. Fractions of dauer formation are shown on the y-axis.

The *srg-37(ean179)* deletion removes 31 amino acids surrounding the pocket structure of the G protein-coupled receptor and causes a frameshift mutation for the 46 C-terminal amino acids, together removing 23% (77/324) of the predicted SRG-37 amino acid sequence. Thus, this deletion likely impairs SRG-37 function, which could cause lower ascr#5 sensitivity. We hypothesized that, if *srg-37(ean179)* causes loss of gene function, removal of additional *srg-37* coding sequence would not further reduce the ascr#5 sensitivity of *srg-37(ean179)* wild strains. Using CRISPR-Cas9 genome-editing^24, 25^, we removed most of the coding sequences of *srg-37* from wild strains with both wild-type (reference-like) *srg-37* and the natural *srg-37* deletion (Fig. 3a). Indeed, we observed that a large deletion in *srg-37* did not change the ascr#5 sensitivities of two wild isolates with the natural deletion, but reduced the ascr#5 sensitivities of five wild isolates with reference-like *srg-37* (Fig. 3c, Supplementary Fig. 6), indicating that the natural deletion is likely a loss-of-function allele. Taken together, these results show that deletion of an ascr#5 receptor gene underlies natural variation in the dauer-pheromone response across the *C. elegans* population.

### Selection has shaped the genetic variation of the two duplicated *C. elegans* ascr#5-receptor genes

We performed population genetic analysis across the *srg-36* and *srg-37* region by analyzing the genome sequences of 249 wild strains. Natural selection and demographic change can shift the allele frequency spectrum from neutrality, as measured by Tajima’s D^26^. Purifying selection, a selective sweep, or a recent population expansion can cause accumulation of rare alleles at a given locus, indicated by a negative Tajima’s D value. We found that the Tajima’s D values were lowest across the promoter and coding regions of *srg-36* and increase back to background neutrality rates in the promoter region of *srg-37* (Fig. 3d). To specifically dissect selection features of *srg-36* and *srg-37*, we further analyzed the ratio between the non-synonymous substitution rate (Ka) and the synonymous substitution rate (Ks) of the two genes by comparing nine homologs across four *Caenorhabditis* species^27^. We found that the non-synonymous substitution rate is much lower than the synonymous substitution rate (Ka/Ks < 1) for both genes, and the Ka/Ks statistic of *srg-36* (0.3145) is lower than *srg-37* (0.4223). Differences in deletion allele frequencies between *srg-36* and *srg-37* also correlate with stronger purifying selection at *srg-36*. The 411-bp deletion allele, *srg-36(ean178)*, is only found in a single wild isolate (PB303), whereas 18.4% (46/249) of wild strains carry the 94-bp deletion allele, *srg-37(ean179)*. These results together suggest that the two pheromone receptor genes are both under purifying selection, but the selection pressure is stronger for *srg-36*.

Although *srg-36* and *srg-37* are duplicated genes that are specific to the same ligand and expressed in the same cells, differences in non-coding and coding sequences between the two genes can cause differences in gene expression levels and receptor activities. Notably, previous studies report that transgene expression of *srg-36* showed a stronger effect than *srg-37* on the ascr#5 response^19^. To test whether *srg-36*, which is likely under stronger purifying selection than *srg-37*, plays a larger role in the ascr#5 response, we performed loss-of-function experiments. We removed the entire *srg-36* coding region in two wild strains – JU346 with wild-type (reference-like) *srg-37* and NIC166 with the natural *srg-37* deletion (Supplementary Fig. 4). First, we found that *srg-36(lf)* reduced ascr#5 sensitivity of both strains, indicating that *srg-36* is functional in both genetic backgrounds (Fig. 3e). Second, we observed that loss of *srg-36* reduced ascr#5 sensitivity more than loss of *srg-37*, supporting the conclusion that *srg-36* plays a larger role than *srg-37* in the ascr#5 response.

The higher activity of *srg-36* could be explained by differences in gene expression levels. We investigated the relative levels of *srg-36* and *srg-37* at the L1 stage, when these genes play critical roles in the dauer-pheromone response, and found that the expression levels of both genes are not significantly different^28^. It is more likely that differences in protein-coding sequences cause the functional differences in the ascr#5 response. Although SRG-36 and SRG-37 show similarities in size and transmembrane structures (Supplementary Fig. 7)^29^, only 46.4% of the amino acid residues are conserved between both receptors. The molecular differences between the two ascr#5 receptors could cause quantitative differences in ascr#5-receptor activities. Taken together, we hypothesized that *srg-36* is the primary ascr#5-receptor gene and is maintained across the *C. elegans* species through purifying selection. By contrast, the redundancy of these two genes facilitates *srg-37* variation and a loss-of-function allele can arise and spread across the population.

### The *srg-37* deletion has spread globally and outcrossed with diverse genotypes

We investigated the locations where wild strains with the natural *srg-37* deletion were isolated and found 46 wild strains with this allele were isolated from all six continents (Fig. 4a, b). Given the low probability of acquiring the same 94-bp deletion, we hypothesized that this allele did not independently arise across multiple global locations but originated from a single ancestral population and spread throughout the world. To test this hypothesis, we analyzed the haplotype composition of *C. elegans* wild isolates across the X chromosome. We reproduced previous studies that showed a recent global selective sweep on the X chromosome^30^, and the shared haplotype on the X chromosome we call the swept haplotype here (Supplementary Fig. 8, 9). Notably, we found that all 46 strains with the *srg-37* deletion exclusively share the swept haplotype at the *srg-37* locus (Fig. 4c). This result not only demonstrates that this allele arose at a single location, but also implies that it has spread throughout the world along with the recent selective sweep. Because the *srg-37* locus is far from the most swept part of the X chromosome, many strains must have outcrossed, suggesting that *srg-37* is unlikely the driver of the X chromosome sweep. Specifically, we found that 34.1% (85/249) of wild strains have an X chromosome that is swept more than 50% of its length but have diverse non-swept haplotypes at the *srg-37* locus (Supplementary Fig. 10). These results suggest that wild strains have undergone outcrossing of the *srg-37* locus, which in turn purged the *srg-37* deletion from these swept subpopulations (Fig. 4d). Taken together, the results suggest that the *srg-37* deletion spread globally with the selective sweeps but was lost because of more recent outcrossing.

**Fig. 4:**
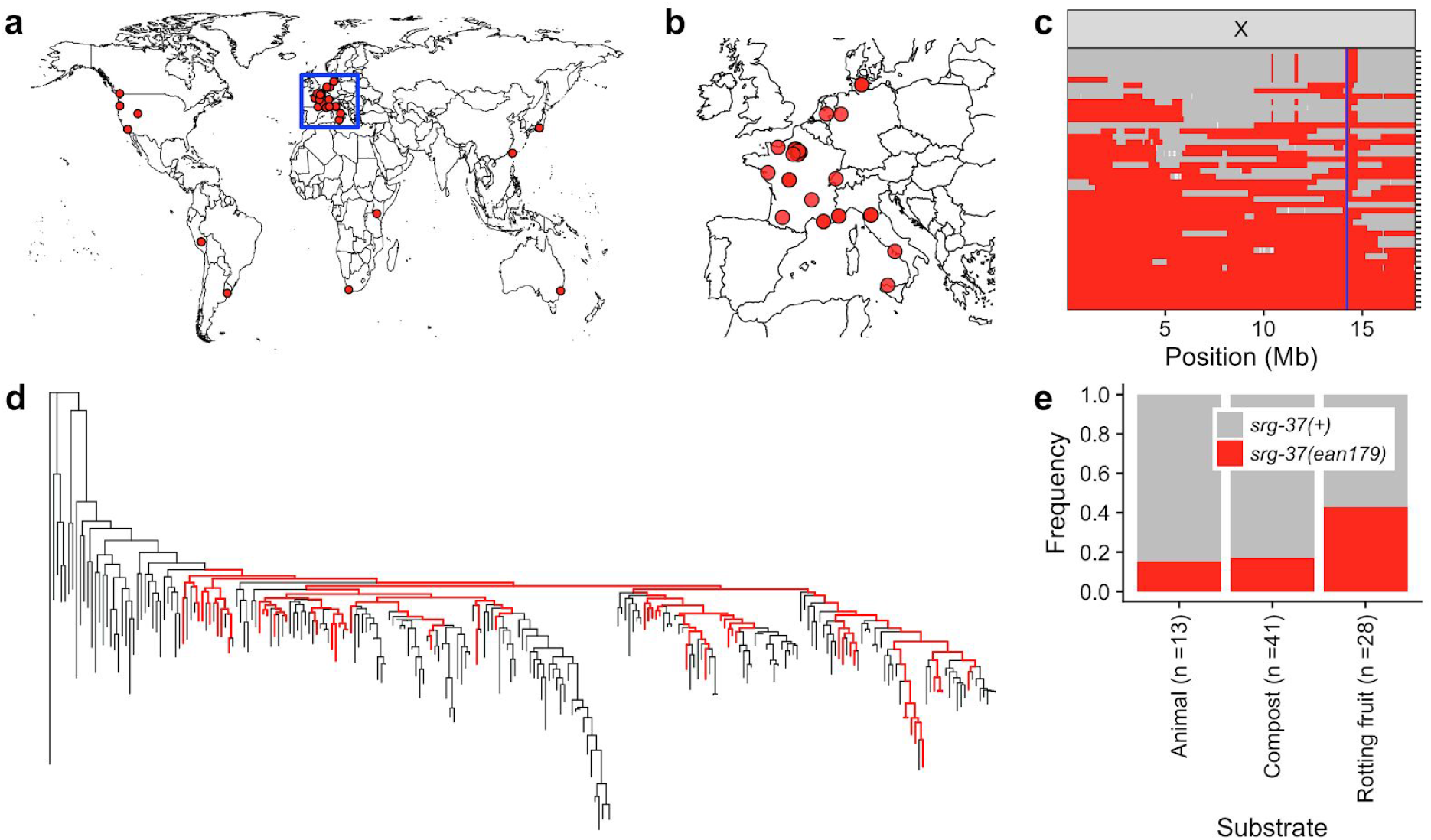
Worldwide and niche-associated gene flow shape the ascaroside (ascr#5) pheromone receptor locus. (a) The global distribution of wild strains that contain the *srg-37(ean179)* deletion allele (red circle) is shown. (b) The geographic distribution of wild strains that are sampled from Europe (inset). Wild strains that contain the *srg-37* deletion (red circle) are shown. (c) Sharing of the swept haplotype on the X chromosome among 46 wild isolates with *srg-37(ean179)* is shown. Each row is one of the 46 strains, ordered roughly by the extent of swept-haplotype sharing (red). Other haplotypes are colored grey. Genomic position of X chromosome is shown on the x-axis. The blue line shows the position of the *srg-37* locus. (d) The genome-wide tree of 249 *C. elegans* wild isolates with those strains that have the *srg-37* deletion shown as red. (e) Stacked bar plots of *srg-37(+)* (grey) and *srg-37(ean179)* (red) allele frequencies among three subpopulations that were sampled from different substrates across hybrid zone in Europe (see Materials and methods). Substrate types and sample sizes are shown on the x-axis, and allele frequencies of each allele are shown on the y-axis. The *srg-37(ean179)* deletion allele is significantly enriched in the rotting fruit subpopulation (*p* = 0.0026).

### The *srg-37* deletion is enriched in the rotting fruit niche

Dauer development is a trade-off between long-term survival and short-term reproductive success. Because reduction of the dauer-pheromone response can promote reproductive growth, we investigated whether wild strains with the *srg-37* deletion were sampled more often from substrates with proliferating populations. These populations are often found in nutritious habitats, such as rotting vegetation^31^. By contrast, *C. elegans* were sampled predominantly in the dauer stage from animal and compost substrates^6, 32^. We analyzed the allele frequencies of the *srg-37* deletion among three subpopulations sampled from animals, compost, and rotting fruits across geographic locations where both *srg-37* alleles were isolated (Supplementary Fig. 11, see Materials and methods). We found that wild strains with the *srg-37* deletion were 67% enriched in rotting fruits (Fig. 4e, Supplementary Table 1). Thus, this allele is not only associated with lower dauer-pheromone responses but also with natural substrates that are known to support reproductive growth.

## Discussion

Dauer pheromones are chemical signals that are perceived by sensory neurons through the combined actions of chemoreceptors and cGMP-mediated signaling^5, 33^. In the absence of dauer-pheromone signaling, the insulin/IGF-1 and TGF-β signaling pathways promote reproductive growth through the production of steroid hormones (dafachronic acid)^34^. Genetic variation in the genes that mediate pheromone perception or downstream signaling likely alter an individual’s dauer-pheromone response. However, because the signaling pathways that act downstream of pheromone perception are involved in various biological processes^35, 36^, mutations in these pathways might cause deleterious pleiotropic effects. Previous studies have shown that the ascr#5-receptor SRG-37 was lost in two independent laboratory lineages of *C. elegans*^19^, suggesting that selection more readily acts at the pheromone perception step of this developmental pathway. In this study, we provide further support for this hypothesis by showing that 18% of wild *C. elegans* strains harbor a putative loss-of-function deletion in the ascr#5-receptor SRG-37, and that these individuals are more likely to be found in nutrient-rich habitats. Thus, modification of pheromone-receptor activity might be favored in both laboratory and natural conditions to fine-tune dauer-pheromone responses with few pleiotropic effects^19, 37^. However, we identified additional dauer-pheromone response QTL, suggesting that multiple loci are involved in ascr#5 responses. Interestingly, SRG-36 and SRG-37 are the only two known ascr#5 receptors involved in dauer-pheromone signaling. Therefore, the other three ascr#5-response QTL indicate that additional natural genetic variants could affect uncharacterized ascr#5 receptors, novel or known factors that regulate receptor activity, or downstream signaling components.

Insights into the redundant functions of *srg-36* and *srg-37* were first gained from the observation that both genes were deleted from two independent laboratory-domesticated *C. elegans* lineages^19^. We did not find a single wild strain in the *C. elegans* population that carries a deletion of both *srg-36* and *srg-37*. Investigations of within (Tajima’s *D*) and among (Ka/Ks) species neutrality statistics suggest that selection acts on these two genes differently. Our results imply that the *srg-36* and *srg-37* genes might not be functionally equivalent in the wild population. The loss-of-function experiments suggest that *srg-36* plays a larger role in the ascr#5 response than *srg-37*. Given the important role of the dauer stage in the long-term survival and the phoretic dispersal of the species, purifying selection might act to conserve the primary ascr#5 receptor (SRG-36) in the *C. elegans* population to maintain the dauer-pheromone response.

The gene-dosage hypothesis proposes that duplicated genes can each become fixed in a population if a dosage increase causes a selective advantage^38^. Recent studies in the natural budding yeast population have reported that copy number variants (CNVs) can explain larger proportions of trait variance compared to single-nucleotide variants (SNVs)^39^. It was also shown that a large fraction of human olfactory-receptor genes exhibit copy number variation^40–43^, so signal perception might be a fruitful substrate for trait evolution. We show that dosage differences of the ascr#5 receptor genes explain the largest dauer-pheromone response QTL, suggesting a pervasive effect of CNVs across species. Our observation that the *srg-37* deletion is enriched in rotting fruits provides evidence that selection can act on CNVs in a niche-specific manner. Although our studies focus on the ascr#5 receptors, approximately 1,300 chemoreceptor genes and 400 pseudogenes are annotated in the reference *C. elegans* genome^44, 45^. By contrast, parasitic nematodes tend to have fewer chemoreceptors^46^, and these species inhabit less heterogeneous niches. Therefore, we hypothesize that the expanded diversity of chemoreceptors and the prevalence of chemoreceptor pseudogenes in free-living *Caenorhabditis* species is caused by broad context-dependent selection in diverse niches. Furthermore, the diversity of the chemoreceptor repertoires is captured across wild *C. elegans* strains that are sampled from various niches^47, 48^. Taken together, these observations indicate that niche adaptation could facilitate the dynamic duplication, deletion, and functional diversification of chemoreceptor gene evolution^49, 50^.

## Materials and methods

### C. elegans strains

Animals were cultured at 20°C on modified nematode growth medium (NGMA) seeded with the *E. coli* strain OP50^51^. Prior to each assay, strains were passaged for at least four generations without entering starvation or encountering dauer-inducing conditions. For the genome-wide association (GWA) studies, 157 wild isolates from CeNDR (version 20170531) were used^22, 23^. All strain information can be found in Supplementary Table 2.

### High-throughput dauer assay

Strains were propagated for four generations on agar plates, followed by bleach synchronization. Approximately 50 embryos were titered and placed into each well of a 96-well microtiter plate filled with 50 μL of K medium^52^ with modified salt concentrations (10.2 mM NaCl, 32 mM KCl, 3 mM CaCl_2_, 3 mM MgSO_4_), 50 μM kanamycin, 5 mg/mL HB101 bacterial lysate (Pennsylvania State University Shared Fermentation Facility, State College, PA), and synthetic ascaroside^53^ dissolved in 0.4% ethanol or 0.4% ethanol alone. Animals were cultured for 52 hours at 25°C until they reached the young adult stage or arrested at the dauer stage. Animals were exposed to 0.5 μm fluorescent microspheres (Polysciences, cat. # 19507-5) at a final concentration of 7.28 × 10^8^ particles/mL and 5 μL of 1 mg/mL HB101 bacterial lysate to promote feeding for 20 minutes. After this exposure, 200 µL of 50 mM sodium azide was added to each well to kill the animals, stop feeding, and straighten the animals. Using the COPAS BIOSORT large particle flow cytometer (Union Biometrica, Holliston MA), optical parameters of animals, including fluorescence intensity, time-of-flight (TOF, animal length), and extinction (optical density) were measured. Measured parameters were used to build a model that can differentiate dauer and adult stages of the population in each well through the R package EMCluster^54^. One cluster with lower fluorescence and smaller body size was assigned to the dauer population and the other to the non-dauer population. The dauer fraction was calculated per well as a fraction of dauer animals among total animals, which is shown as a single data point in each plot. From the control experiments, both the false positive ratio (false dauer detection in a wild-type sample without pheromone treatment) and the false negative ratio (false non-dauer detection in Daf-c mutant sample) were 5%, indicating 95% accuracy of the assay (Fig. 1b,c).

### Heritability calculations

For dose-response experiments, broad-sense heritability (*H*^2^) estimates were calculated using the *lmer* function in the lme4 package with the following linear mixed model (phenotype ∼1 + (1|strain))^55^. *H*^2^ was then calculated as the fraction of the total variance that can be explained by the random component (strain) of the mixed model.

### Genome-wide association mapping

A genome-wide association (GWA) mapping was performed using phenotype data from 157 wild *C. elegans* strains. Genotype data were acquired from the latest VCF release (Release 20180527) from CeNDR that was imputed as described previously^22^. We used BCFtools^56^ to filter variants that had any missing genotype calls and variants that were below 5% minor allele frequency. We used PLINK v1.9^57, 58^ to LD-prune the genotypes at a threshold of *r*^2^ < 0.8, using --*indep-pairwise 50 10 0.8*. The pruned genotype set comprised 72,568 markers that were used to generate the realized additive kinship matrix using the *A.mat* function in the *rrBLUP* R package^59^. These markers were also used for genome-wide mapping. However, because these markers still have substantial LD within this genotype set, we performed eigen decomposition of the correlation matrix of the genotype matrix using *eigs_sym* function in Rspectra package^60^. The correlation matrix was generated using the *cor* function in the correlateR R package^61^. We set any eigenvalue greater than one from this analysis to one and summed all of the resulting eigenvalues^62^. This number was 915.621, which corresponds to the number of independent tests within the genotype matrix. We used the *GWAS* function in the rrBLUP package to perform genome-wide mapping with the following command: *rrBLUP::GWAS(pheno = dauer, geno = Pruned_Markers, K = KINSHIP, min.MAF = 0.05, n.core = 1, P3D = FALSE, plot = FALSE)*. Regions of interest are defined as +/- 100 SNVs from the rightmost and leftmost markers above the eigen-decomposition significance threshold. If regions of interest for separate QTL are within 1000 SNVs, they become grouped as a single region of interest.

### Identification of natural deletion variants of *srg-36* and *srg-37*

Whole-genome sequence data were aligned to WS245 using bwa (version 0.7.8-r455). Optical/PCR duplicates were marked with PICARD (version 1.111)^22, 63–65^. Alignments with greater than 100X coverage were subsampled to 100X using sambamba^66^. We called large deletions using the Manta structural variant caller (v1.4.0) using the default caller and filter settings^67^.

### Generation of *srg-36* and *srg-37* deletion strains

*srg-36* and *srg-37* loss-of-function mutant strains were generated by CRISPR-Cas9-mediated genome editing, using a co-CRISPR approach and Cas9 ribonucleoprotein (RNP) delivery^24, 25^. crRNAs synthesized by IDT (Skokie, IL) targeting *srg-36* (exon 1 and the 3’ UTR) and *srg-37* (exon 2 and exon 5) were used to generate deletions. The injection mixture (10 μL) was prepared with 0.88 μL of 200 μM tracrRNA (IDT, Product #1072532), 0.88 μL of 100 μM crRNA1 (5’ targeting) and crRNA2 (3’ targeting), and 0.12 μL of 100 μM *dpy-10* crRNA (IDT) were mixed and incubated at 95°C for five minutes. After cooling to room temperature, 2.87 μL of 60 μM Cas9 protein (IDT Product #1074181) was added and incubated at room temperature for five minutes. Finally, 0.5 μL of 10 μM *dpy-10* ssODN (IDT) repair template and 3.99 μL of nuclease-free water were added. RNP injection mixtures were microinjected into the germline of young adult hermaphrodites (P0), and injected animals were singled to fresh 6 cm NGM plates 18 hours after injection. Two days later, F1 progeny were screened, and animals expressing a Rol phenotype were transferred to new plates and allowed to generate progeny (F2). Then, F1 animals were genotyped by PCR. Deletion of *srg-36* was detected with primers oECA1460-1463 and deletion of *srg-37* was detected with primers oECA1429, oECA1430, and oECA1435. Non-Rol progeny (F2) of F1 animals positive for the desired deletion were propagated on separate plates to generate homozygous progeny. F2 animals were genotyped afterwards with same primer sets, and PCR products were Sanger sequenced for verification. All crRNA and oligonucleotide sequences are listed in the Supplementary Table 3.

### Population genetics

Sliding window analysis of population genetic statistics was performed using the PopGenome package in R^68^. All sliding window analyses were performed using the imputed SNV VCF available on the CeNDR website with the most diverged isotype XZ1516 set as the outgroup^22, 69, 70^. Estimates of Ka/Ks were performed using an online service (http://services.cbu.uib.no/tools/kaks). Linkage disequilibrium (LD) of QTL markers, which can be measured as the square of the correlation coefficient (*r^2^*), was calculated using the genetics package in R^71^. The formula for the correlation coefficient is *r = −D / sqrt (p(A) * p(a) * p(B) * p(b))*. Haplotype composition of each wild isolate was inferred by applying IBDseq^72^ with variants called by BCFtools^73^ and the following filters: Depth (DP) > 10; Mapping Quality (MQ) > 40; Variant quality (QUAL) > 10; (Alternate-allelic Depth (AD) / Total Depth (DP)) ratio > 0.5; < 10% missing genotypes; < 10% heterozygosity. To generate genome-wide tree, whole-population relatedness analysis was performed using RAxML-ng with the GTR+FO substitution model(DOI:10.5281/zenodo.593079). SNVs were LD-pruned using PLINK (v1.9) with the --indep-pairwise command ‘--indep-pairwise 50 1 0.95’. We used the vcf2phylip.py script (DOI:10.5281/zenodo.1257058) to convert the pruned VCF files to the PHYLIP format^74^ required to run RAxML-ng. To construct the tree that included 249 strains, we used the GTR evolutionary model available in RAxML-ng^75, 76^. Trees were visualized using the ggtree (v1.10.5) R package^77^.

### Substrate specificity analysis in the co-sampling zone

The co-sampling zone was defined as a location where both *srg-37(+)* and *srg-37(ean179)* were isolated (Supplementary Fig. 11). Collection information available on the CeNDR website were used to analyze correlation between isolated substrate and the *srg-37* genotype of each isolate. Isolation of wild strains that share same genome-wide genotypes (isotype) were counted as independent isolations if they were sampled from different locations or from different substrate types. We found that 95 isotypes were isolated in co-sampling zone from at least 119 independent isolations. Three substrates (animals, compost, rotting fruit) with more than ten independent isolated strains were selected for substrate enrichment test. In total, 82 wild strains (66 isotypes) were grouped into three subpopulations by the substrate where they were isolated, and allele frequencies of each subpopulation were calculated. Significant enrichment of *srg-37(ean179)* in each subpopulation was determined by hypergeometric tests using stats R package^78^.

## Acknowledgments

This work was supported by an NSF CAREER Award to E.C.A. S.Z was supported by the Cell and Molecular Basis of Disease training grant (T32GM008061) and The Bernard and Martha Rappaport Fellowship. D.E.C received the National Science Foundation Graduate Research Fellowship (DGE-1324585). Members of the Andersen Lab helped manuscript editing and Ying K. Zhang for assistance with the synthesis of ascarosides. Some strains were provided by the CGC, which is funded by the NIH Office of Research Infrastructure Programs (P40 OD010440). We want to thank WormBase for providing genome data of *Caenorhabditis* species.

## Author contributions

D.L. and E.C.A. conceived and designed the study. D.L. performed the high-throughput assay, CRISPR-Cas9 genome-editing, population genomic analyses, and niche enrichment tests. S.Z. performed the GWA mapping, identified genetic variants in the *dauf-1* locus, generated the genome-wide tree of 249 wild *C. elegans* strains, and edited the manuscript. D.E.C. analyzed the haplotype composition of 249 wild strains. L.F., J-C.H., M.G.S., J.A.G.R., J.W., J.E.K., C.B., and M-A.F. contributed wild isolates to the 249 wild *C. elegans* strain collection. F.C.S provided the dauer pheromone. D.L. and E.C.A. analyzed the data and wrote the manuscript.

---

**Fig. S1:**
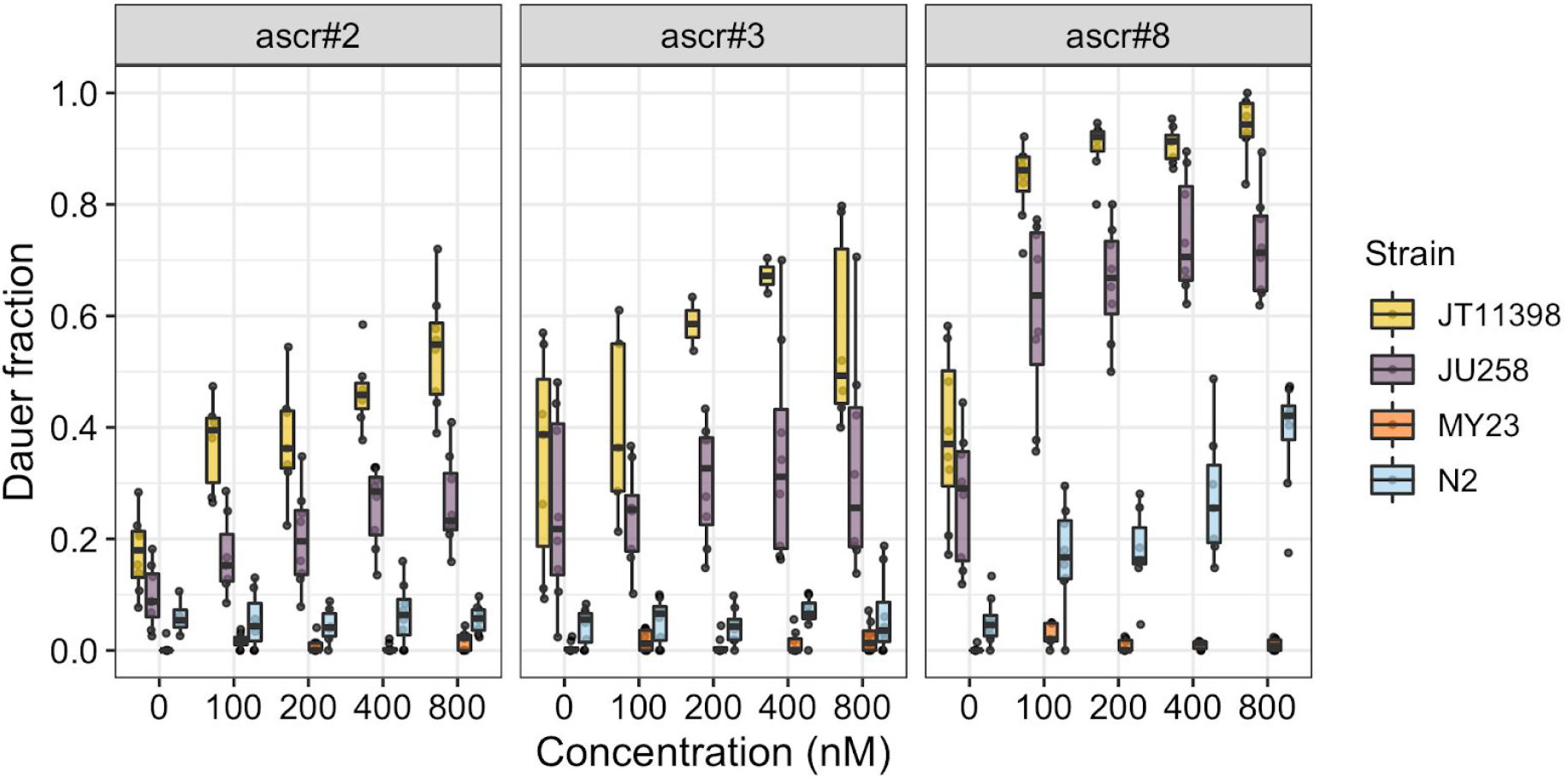
Genetically divergent wild isolates show different dose responses to dauer pheromone (ascr#2, ascr#3, and ascr#8) treatments. Tukey box plots of the dose responses at 25°C for ascr#2 (left), ascr#3 (center), and ascr#8 (right) for four divergent strains are shown with data points plotted behind. Box plots are colored by strain (JT11398 (yellow), JU258 (purple), MY23 (orange) and N2 (skyblue)). Concentrations of ascarosides are shown on the x-axis, and fractions of dauer larvae are shown on the y-axis.

**Fig. S2:**
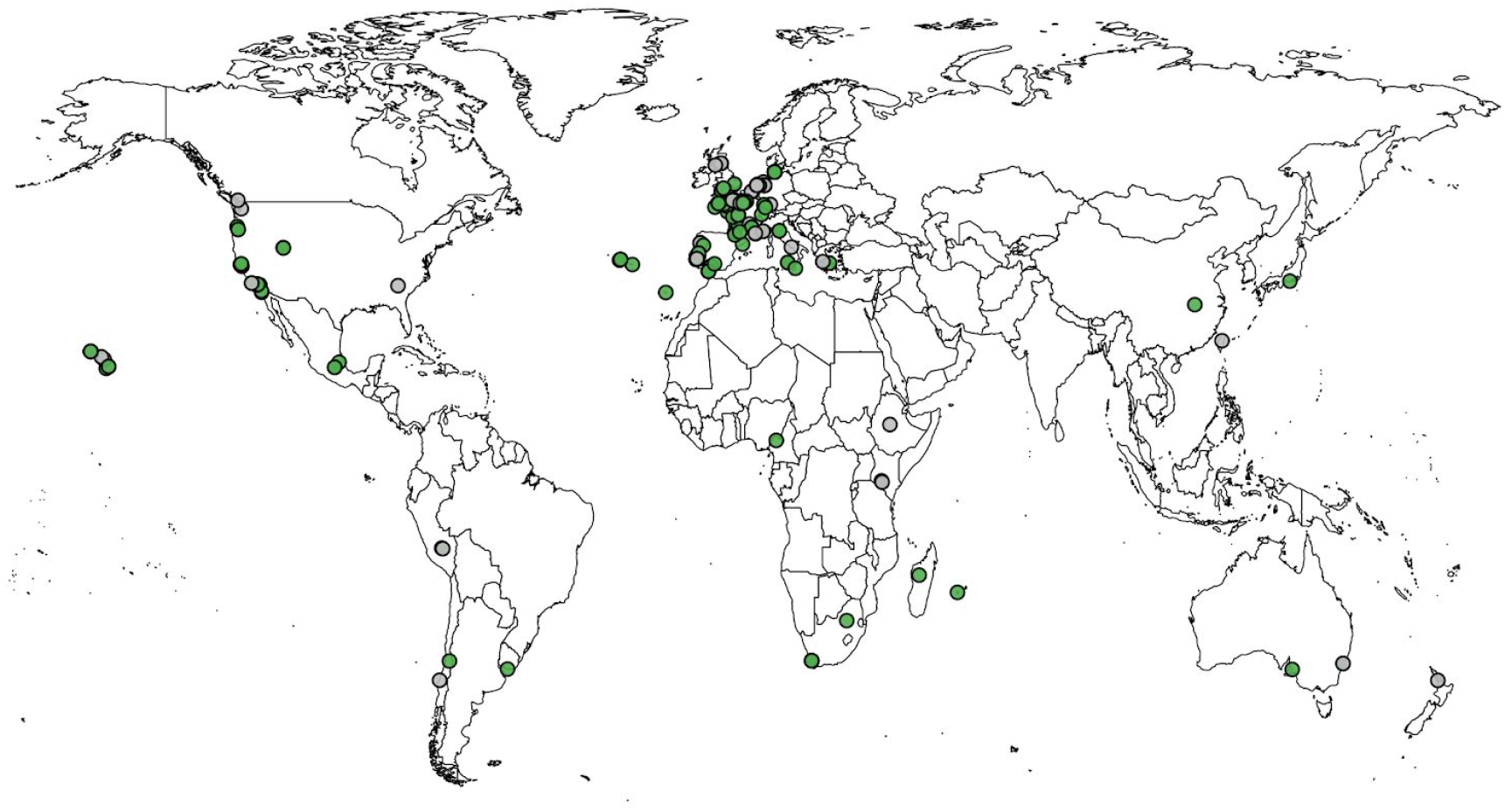
Wild *C. elegans* strains have isolated across six continents. Among the 249 wild strains that are available through the CeNDR, the global distribution of the 239 wild strains with known geographic origins is shown. Among the 157 wild strains that were tested using the HTDA, 151 wild strains with known geographic origins are shown as green circles. Wild strains that are not tested using the HTDA are shown as grey circles.

**Fig. S3:**
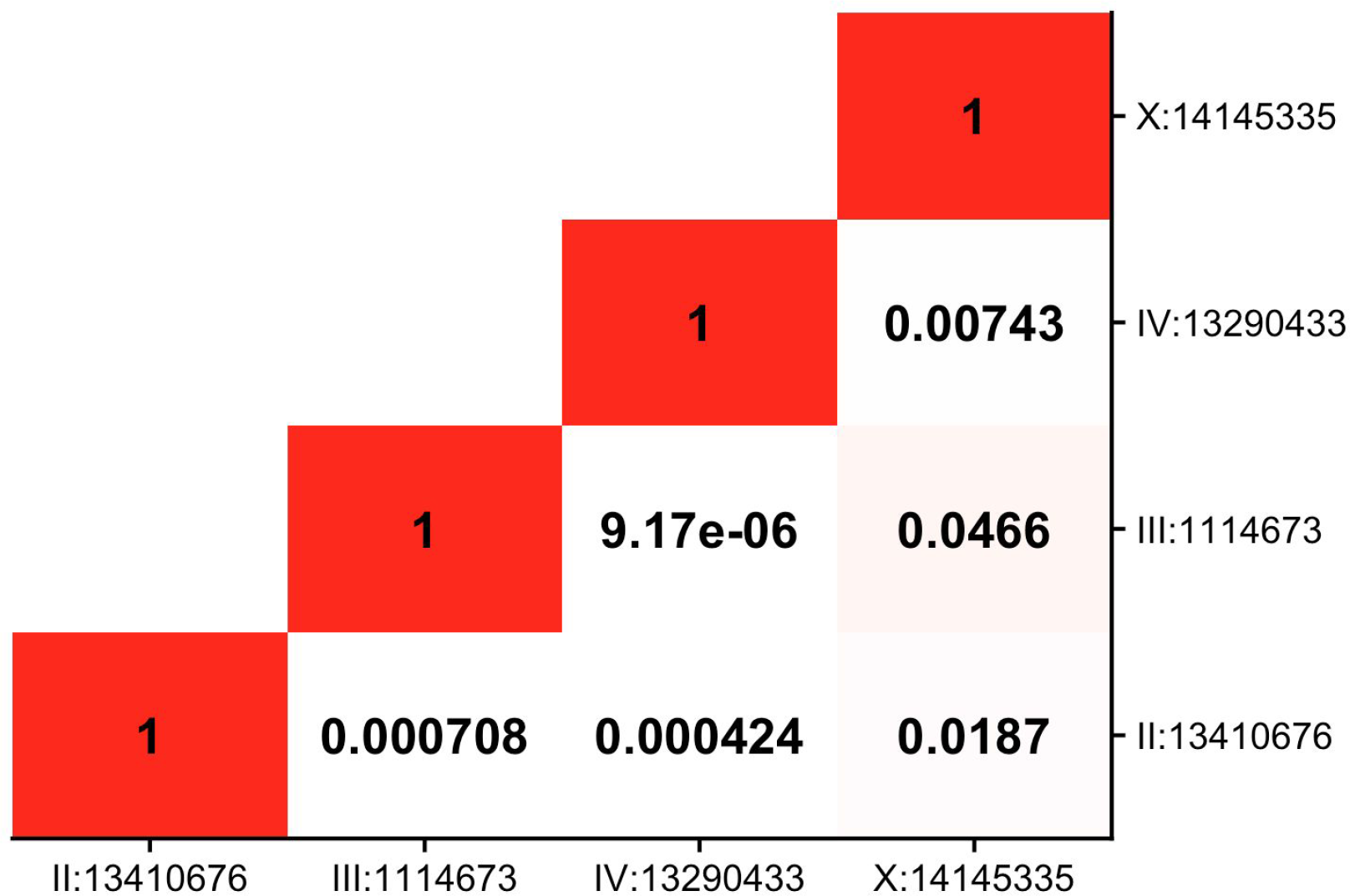
Linkage disequilibrium is not observed among four QTL for ascr#5 response. A heatmap plot that shows linkage disequilibrium (LD) among peak QTL markers as measured by the square of the correlation coefficient (*r^2^*).

**Fig. S4:**
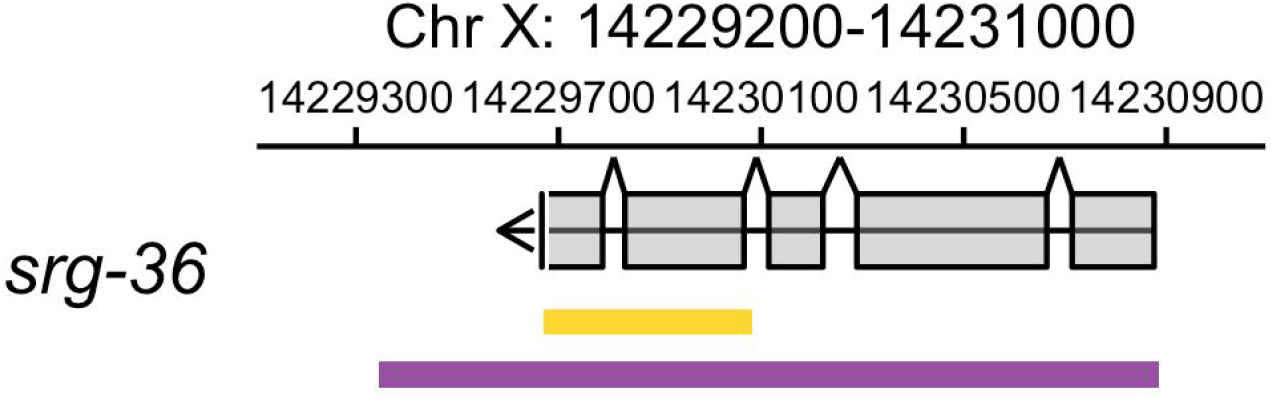
A schematic plot for *srg-36* gene structure. A schematic plot for the *srg-36* gene structure (grey), the 411-bp natural deletion allele *ean178* (yellow), and the CRISPR-Cas9 loss-of-function deletion (purple) are shown.

**Fig. S5:**
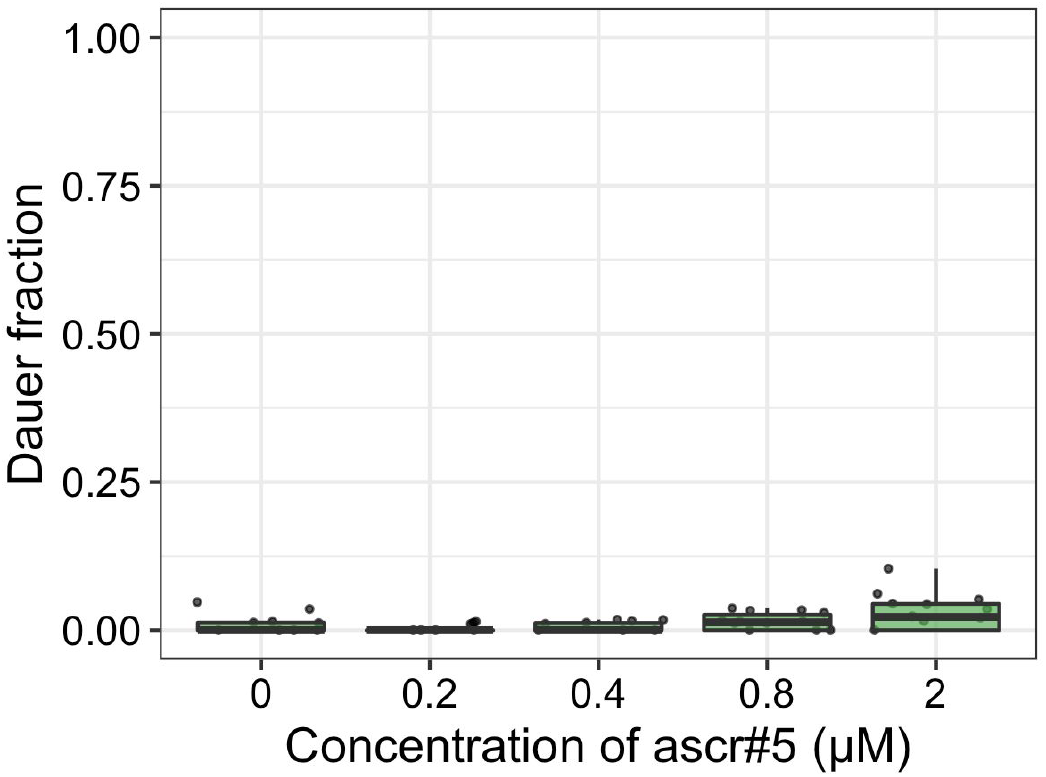
PB303, a unique strain that has a deleted *srg-36*, is insensitive to ascr#5. Tukey box plots of the ascr#5 dose response at 25°C for PB303 that carries the *srg-36(ean178)* deletion allele are shown with data points plotted behind. Concentrations of ascr#5 are shown on the x-axis, and the fraction of dauer formation is shown on the y-axis.

**Fig. S6:**
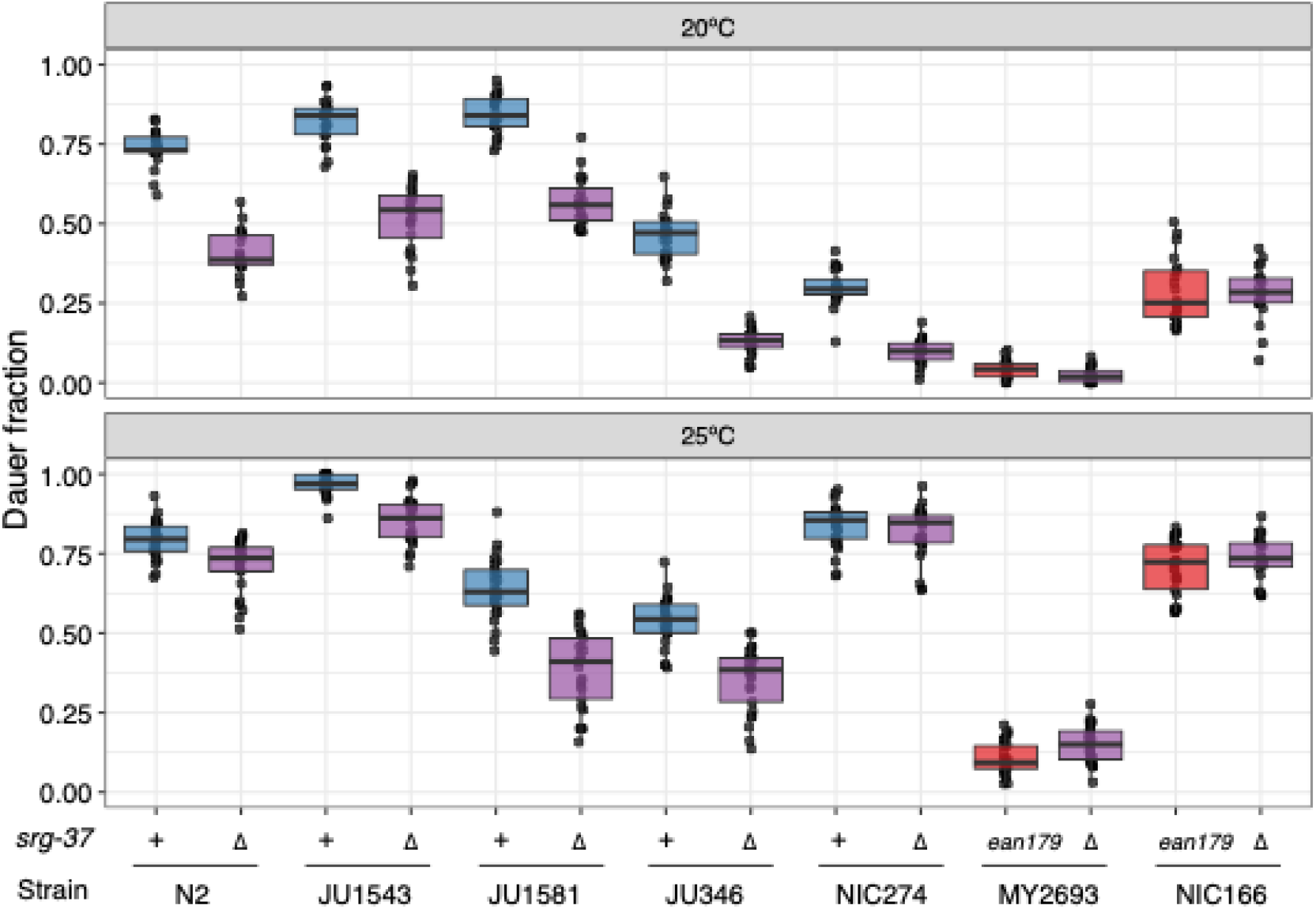
The *srg-37(ean179)* deletion is a putative loss-of-function allele. Tukey box plots of *srg-37* loss-of-function experiments in dauer pheromone conditions (800 nM of ascr#5) at 20°C (top) and 25°C (bottom) are shown with data points plotted behind. Each box plot is colored by the genotypes of *srg-37*, where *srg-37(+)* is blue, *srg-37(ean179)* is red, and CRISPR-Cas9-generated *srg-37(lf)* is purple. Wild isolates and the putative *srg-37* loss-of-function deletion mutants for each genetic background (Δ) are paired on the x-axis. The fraction of dauer formation is shown on the y-axis.

**Fig. S7:**
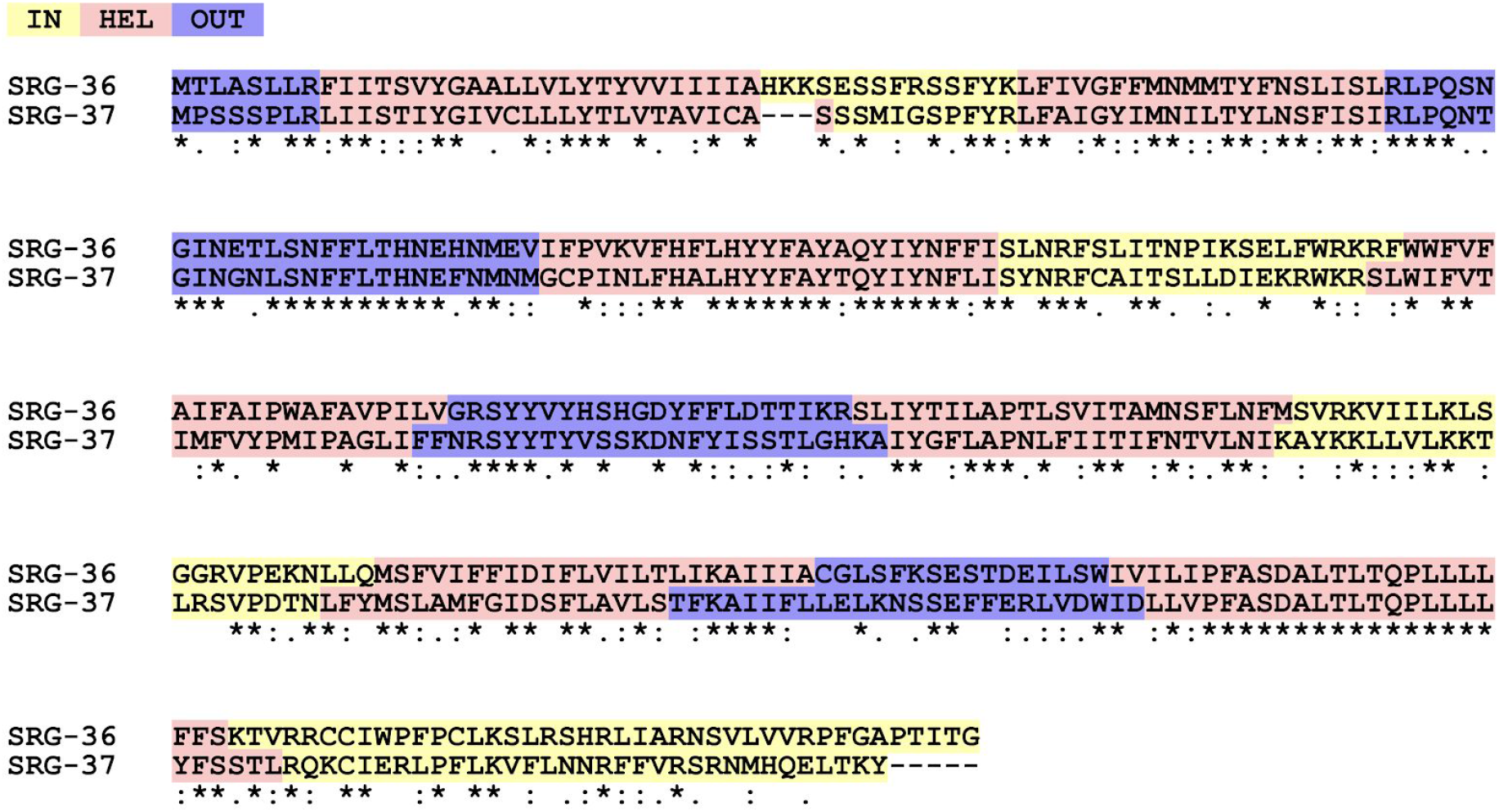
An alignment of SRG-36 and SRG-37 shows conserved transmembrane receptor structure. Reference sequences from the N2 strain were used. Alignment and structure estimations were performed by PSI-TM/Coffee software^29^. Amino acid residues of each protein are colored by putative structural domains (intracellular domains (yellow), transmembrane domain (pink), extracellular domains (blue)). Conservation is annotated below each aligned amino acid (conserved residue (asterisk), strongly conserved residue (> 0.5 in the Gonnet PAM 250 matrix, colon), weakly conserved residues (=< 0.5 in the Gonnet PAM 250 matrix, period).

**Fig. S8:**
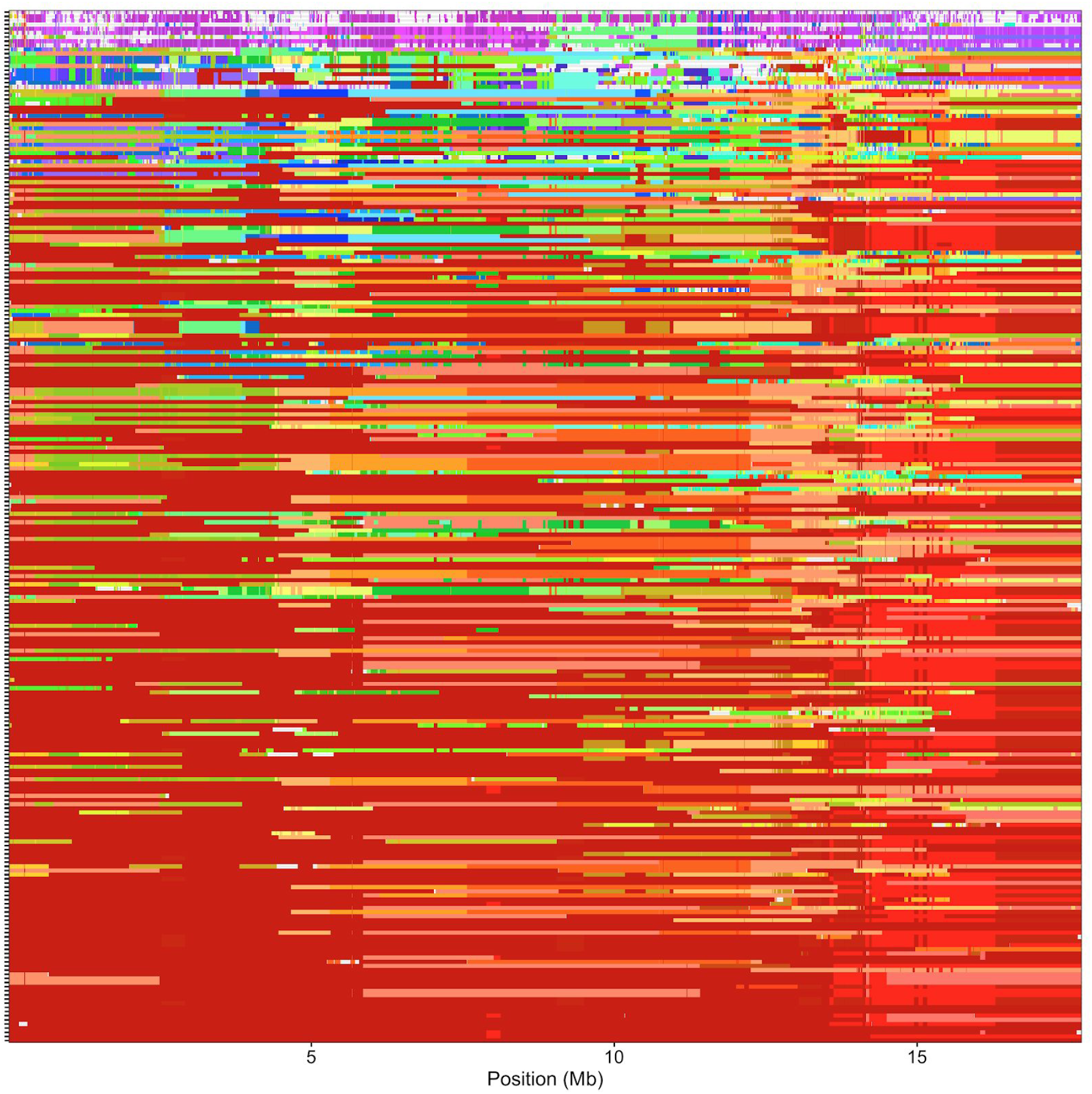
X chromosome sharing of 249 wild strains reveals swept regions where haplotype diversity is reduced. X chromosome sharing plots with color-coded haplotype segments of the 249 wild strains of *C. elegans* are shown. Each row corresponds to genetically unique single wild strains (isotypes), ordered roughly by the extent of haplotype sharing. Shared chromosome regions are shown with the same color. Genomic position on the X chromosomes is shown on the x-axis. Note that similarity in colors does not imply similarity in genotype.

**Fig. S9:**
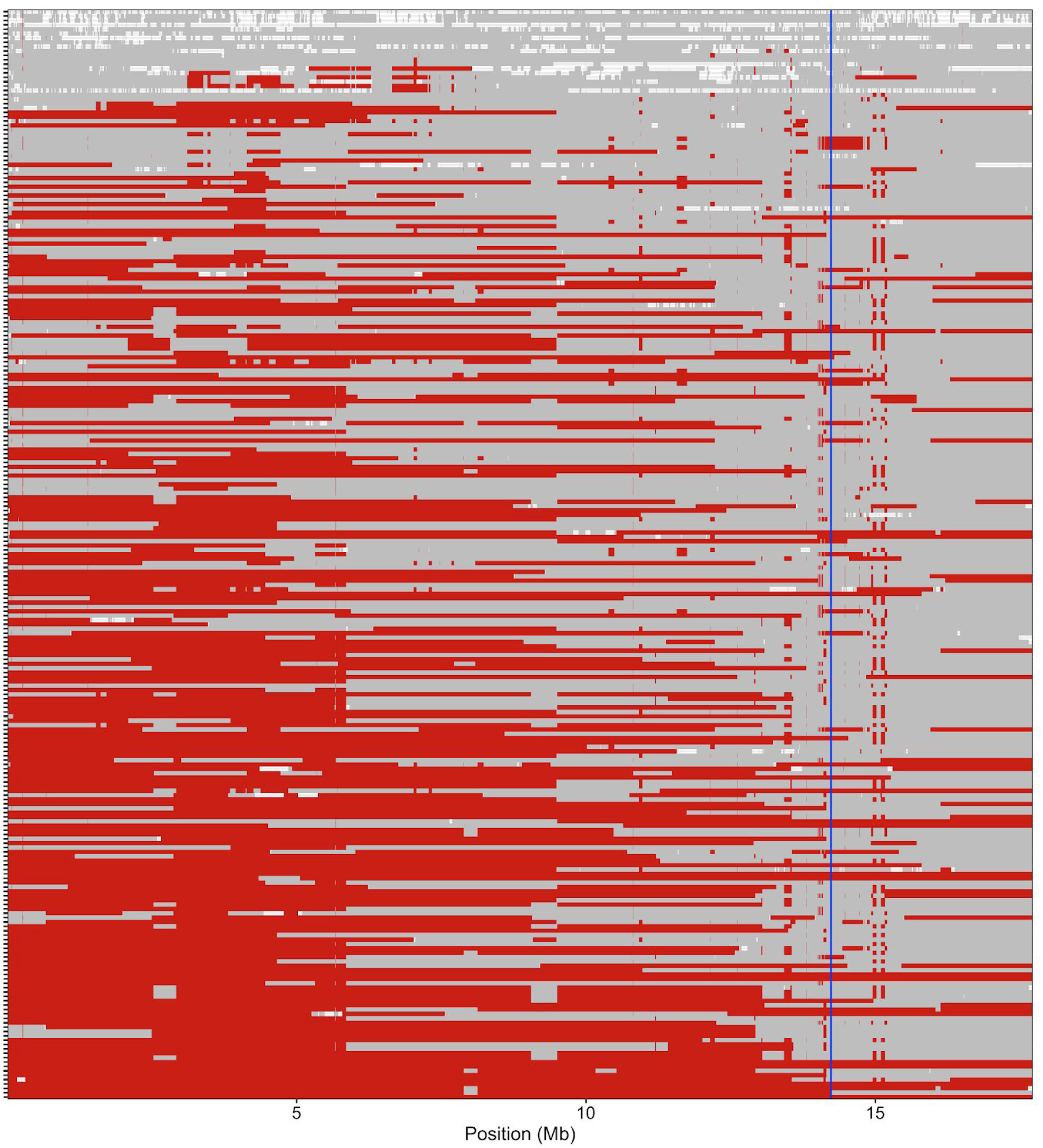
*srg-37* is located far from the left arm of the X chromosome, which is frequently swept across 249 wild *C. elegans* strains. A modified X chromosome sharing plots with the swept haplotype colored as red and other haplotypes as grey are shown. The swept haplotype is defined as a haplotype that is identical-by-descent and most frequently shared across 249 wild *C. elegans* strains. Each row shows the genotype of one of the 249 wild isolates, ordered by the extent of swept haplotype sharing. Genomic position is shown on the x-axis. The blue vertical lines denote the *srg-36 srg-37* locus.

**Fig. S10:**
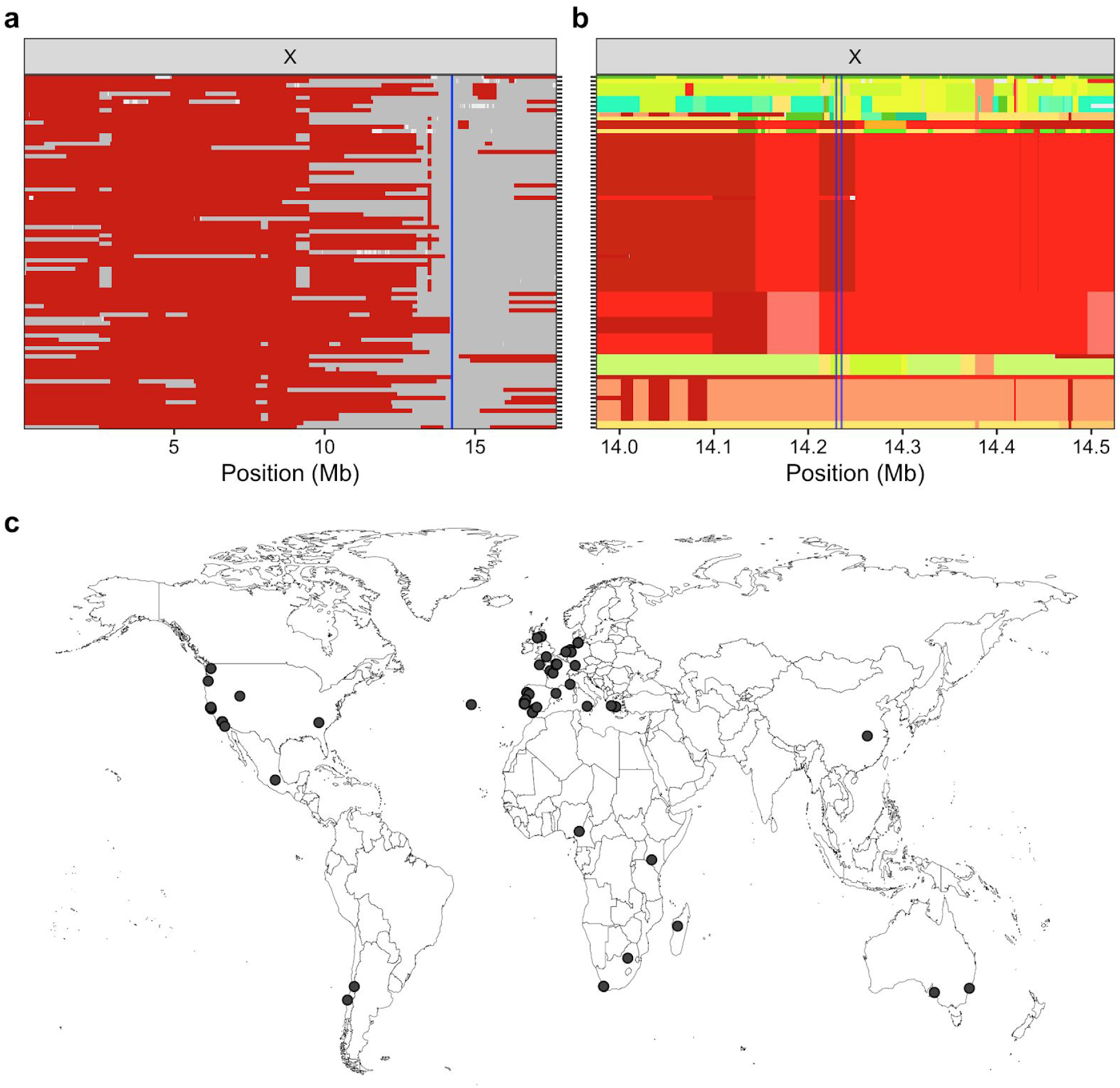
*srg-37(ean179)* is outcrossed by multiple genotypes. The outcrossed subpopulation is defined as a subset of strains with the swept haplotype segments that span more than 50% of the X chromosome except *srg-37* locus. This outcrossed subpopulation comprises 85 wild strains. (a) A modified haplotype sharing plot with the swept haplotype colored as red and other haplotypes colored grey. Each row is one of the wild isolates that belongs to the outcrossed subpopulation, ordered roughly by the extent of the swept haplotype sharing. (b) A haplotype sharing plot with color-coded haplotype segments of outcrossed subpopulation across the X chromosome region (X: 14.0 Mb - 14.5 Mb) flanking and including *srg-36* and *srg-37*. Note that red and dark red color are not swept haplotypes. All shown haplotypes were color-coded by grey in (a). (a, b) The blue vertical lines denote the *srg-36 srg-37* locus. (c) The geographic locations of 85 outcrossed wild strains are shown as grey circles.

**Fig. S11:**
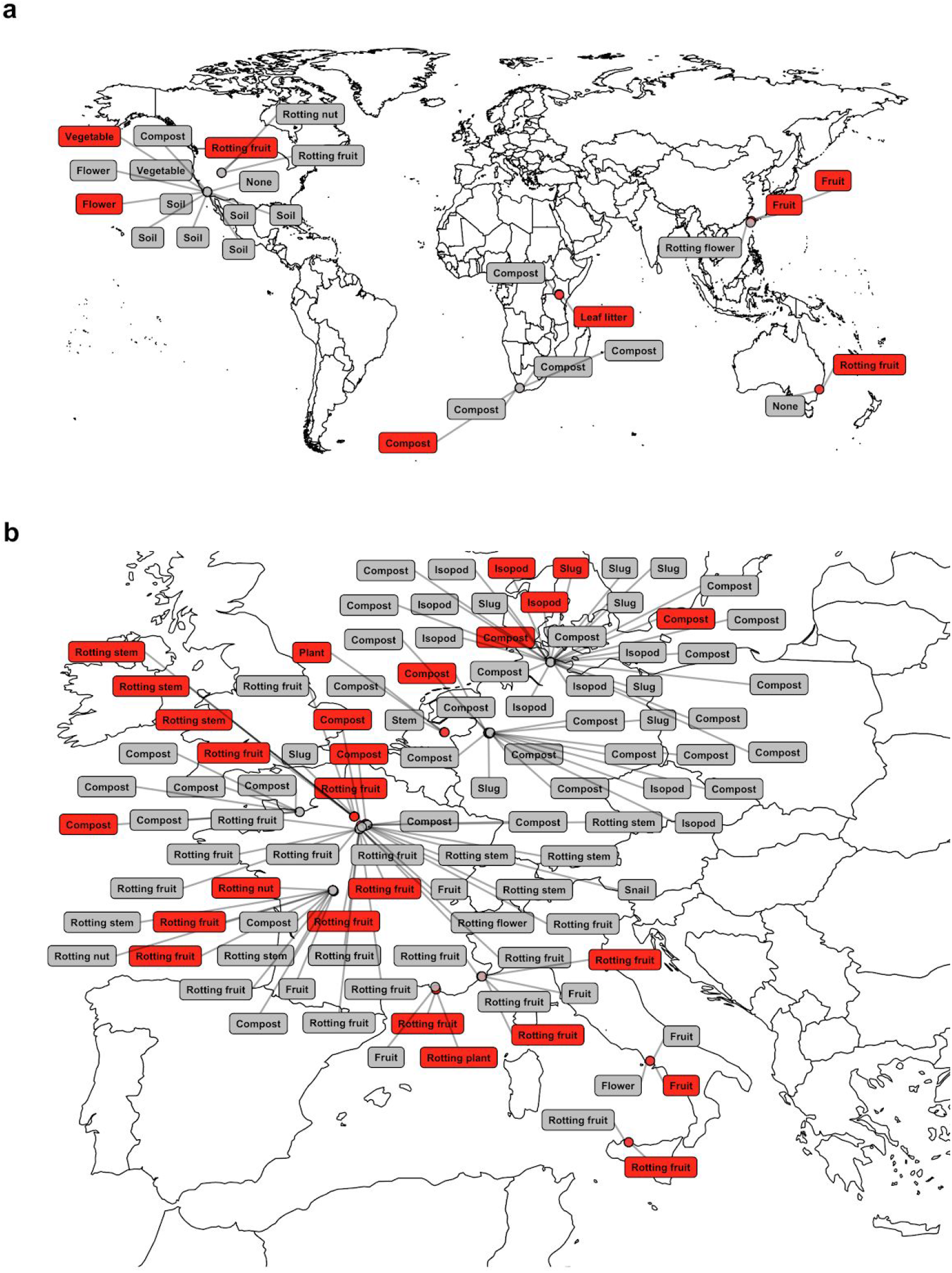
Two *srg-37* genotypes are isolated in the local habitat across the world. Wild strains that are isolated from locations where both genotypes of *srg-37* were found are shown on the map of (a) world and (b) Europe. Each box indicates the substrate type where each strain is isolated, and is colored by the *srg-37* genotype of the isolated strain (grey: wild-type, red: *ean179*).

